# An open-source MRI compatible frame for multimodal presurgical mapping in macaque and capuchin monkeys

**DOI:** 10.1101/2024.02.17.580767

**Authors:** Lucy Liang, Isabela Zimmermann Rollin, Aydin Alikaya, Jonathan C. Ho, Tales Santini, Andreea C. Bostan, Helen N. Schwerdt, William R. Stauffer, Tamer S. Ibrahim, Elvira Pirondini, David J. Schaeffer

## Abstract

**Highlights:**

- We present a compact MRI-compatible stereotaxic frame for large nonhuman primates.
- The design is 3D printable, inexpensive, and matches size of an adult human head.
- Enabled real-time, accurate, MRI-guided deep-brain viral vector injection.
- Facilitated multimodal alignment for deep-brain electrophysiology planning.
- All computer-aided-design files are modularized and publicly available and editable.

**Background:** High-precision neurosurgical targeting in nonhuman primates (NHPs) often requires presurgical anatomy mapping with noninvasive neuroimaging techniques (MRI, CT, PET), allowing for translation of individual anatomical coordinates to surgical stereotaxic apparatus. Given the varied tissue contrasts that these imaging techniques produce, precise alignment of imaging-based coordinates to surgical apparatus can be cumbersome. MRI-compatible stereotaxis with radiopaque fiducial markers offer a straight-forward and reliable solution, but existing commercial options do not fit in conformal head coils that maximize imaging quality.

**New method:** We developed a compact MRI-compatible stereotaxis suitable for a variety of NHP species (*Macaca mulatta*, *Macaca fascicularis*, and *Cebus apella*) that allows multimodal alignment through technique-specific fiducial markers.

**Comparison with existing methods:** With the express purpose of compatibility with clinically available MRI, CT, and PET systems, the frame is no larger than a human head, while allowing for imaging NHPs in the supinated position. This design requires no marker implantation, special software, or additional knowledge other than the operation of a common large animal stereotaxis.

**Results:** We demonstrated the applicability of this 3D-printable apparatus across a diverse set of experiments requiring presurgical planning: 1) We demonstrate the accuracy of the fiducial system through a within-MRI cannula insertion and subcortical injection of viral vectors. 2) We also demonstrated accuracy of multimodal (MRI and CT) alignment and coordinate transfer to guide a surgical robot electrode implantation for deep-brain electrophysiology.

**Conclusions:** The computer-aided design files and engineering drawings are publicly available, with the modular design allowing for low cost and manageable manufacturing.

## 1. Introduction

Because of their phylogenetic proximity to humans, preclinical research in nonhuman primates (NHPs) has been central for translational research (Friedman et al., 2017; Phillips et al., 2014). With highly variable anatomy across the primate lifespan, high-precision neurosurgical targeting in NHPs often requires presurgical mapping of individual anatomy, with noninvasive neuroimaging techniques (MRI, CT, PET) allowing for translation of individual anatomical coordinates to surgical stereotactic (or robotic) apparatus (Miocinovic et al., 2007). Given the varied contrasts that these imaging techniques produce (e.g., soft versus hard tissue), however, precise alignment of imaging-based coordinates to the surgical apparatus can be inefficient. MRI-compatible stereotaxic devices with radiopaque fiducial markers offer a straight-forward and reliable solution to this issue, but existing commercial options do not allow for use in conformal head coils designed to maximize resolution with a high signal-to-noise ratio that is often required for accurate presurgical mapping. Here, we developed a compact MRI-compatible stereotaxis suitable for a variety of NHP species (*Macaca mulatta*, *Macaca fascicularis*, and *Cebus apella* shown here) that allows for multimodal alignment (MRI, CT, PET) through technique-specific fiducial markers.

Stereotaxic frames are a standard piece of surgical equipment used to ensure accuracy in nearly any brain targeting procedure. When paired with imaging, they provide significant benefits such as alignment to a common atlas or overlaying multiple modalities of scans, enabling precise surgical planning (Alvarez-Royo et al., 1991; Fredericks et al., 2020; Frey et al., 2011; Ho et al., 2022; Miocinovic et al., 2007; Saunders et al., 1990). Indeed, there are several commercially available, as well as custom-designed MRI-compatible large animal stereotaxic frames that provide improved targeting accuracy (Chen et al., 2015; Subramanian et al., 2005). However, because of the large dimensions, these frames are often difficult to use with constantly evolving high field scanners and custom coils that maximize resolution with a high signal-to-noise ratio, especially when placing the animal in the sphinx position that is often used for surgery. To obviate these issues, elegant approaches have been developed to create a ‘frameless’ approach to MR guided stereotactic procedures (Bentley et al., 2018; Dubowitz and Scadeng, 2011; Frey et al., 2004; Ohayon and Tsao, 2012; Paiva et al., 2020; Walbridge et al., 2006), but they require implanted hardware, specific software, or specialty in identifying bony landmarks that may not be practical for one-off experiments where the MRI is only used to guide the surgery, rather than the focus of the experiment. By placing the animal in the supine position and conforming the dimensions of the design to that of the human head, our design allows for ease of imaging across modalities and clinically available MRI, CT, and PET platforms.

In an effort to support the ethical considerations referred to as the “three Rs” - Replacement, Reduction and Refinement (Russell and Burch, 1959), international consortia have arisen to make data and apparatus arising from NHP research sharable and openly available (Fox et al., 2021; Messinger et al., 2021; Milham et al., 2022; Schaeffer et al., 2022). In aid of this progress toward democratization of data and the supporting neuroscientific apparatus, we present designs for a compact MRI-compatible stereotactic frame designed for imaging in the supinated position for the express purpose of compatibility with clinically available MRI, CT, and PET systems in which humans are typically imaged while laying on their backs. With the frame being no larger than the gross dimensions of an adult human head, our design serves to resolve long-standing issues of fitment of the stereotactic apparatus within imaging environments. Despite the compact footprint, the design is applicable across a variety of species including *Macaca mulatta*, *Macaca fascicularis*, and *Cebus apella*. No single component is larger than the build plate size of a standard stereolithography printer (e.g., < 400 cm^3^) and thus the frame is relatively easy to manufacture with standard 3D printing. The frame is intended to be used across modalities, with liquid-tight fiducial cavities for modality-specific contrast enhancement (e.g., MRI T1-contrast enhancers, or PET ligands). We confirmed that the frame does not introduce geometric distortion or impact the signal homogeneity by conducting CT and MRI scans of agar MRI phantom with linear internal geometry that approximate the size of a macaque brain. We further demonstrate the use and accuracy of the frame through 1) an MRI-guided real-time microinjection of adeno-associated virus (AAV) with immunohistological reporting, 2) coregistration of MRI and CT images to perform robot-aided neurosurgical implantation of a deep brain electrode to target different subcortical structures, e.g., the hand area of the corticospinal tract (CST) within the internal capsule (IC).

## 2. Materials and Methods

### 2.1 Animals

Two *Macaca mulatta* monkeys (MK-EF: 6 yo, 8.35 kg, male; MK-HS, 7 yo, 12.00 kg, male), four *Macaca fascicularis* monkeys (MK-SC: 4 yo, 5.40 kg, male; MK-SZ: 4 yo, 5.90 kg, male; MK-JC: 6 yo, 7.50 kg, male; MK-OP: 6 yo, 6.00 kg, female), as well as two *Cebus apella* monkeys (a.k.a. capuchin monkeys, MK-JN: 14 yo, 2.93 kg, female; MK-BB: 16 yo, 4.40 kg, male) were involved in this study. Animals were housed in the Division of Laboratory Animal Resources at the University of Pittsburgh. All procedures performed were approved by the University of Pittsburgh Animal Research Protections and Institutional Animal Care and Use Committee (protocol numbers: 22020612, 22017081, 22091621, and 19034358).

### 2.2 Frame Design

All hardware was designed in concert with veterinary technicians to minimize animal discomfort and improve handling efficiency. The concept behind the design of the frame was to allow placement of standard stereotactic fiducial markers - ear bars, eye bars, and a pallet bar (**Fig. 1A**) while having an outer dimension (165×100×190 mm) approximating that of a human head (**Fig. 2A**). With the supine position being standard for human clinical neuroimaging systems, the frame was designed to securely hold the animal in stereotactic position while supine, allowing for simplified placement in clinical imaging systems (**Fig. 1B, 1C**). The computer aided design (CAD) files were designed in SolidWorks (SOLIDWORKS 2021, Dassault Systèmes, Waltham, Massachusetts, USA). Although large format 3D printers will allow for printing of both halves of the frame as one piece, the provided files were designed to allow for modular printing on small format desktop 3D printers (**Fig. 1D**). The ear bars were designed to accept screw-in extensions that allow for ease of ear bar placement, while minimizing physical interaction with imaging hardware once removed (**Fig. 1A**).

**Figure 1.**
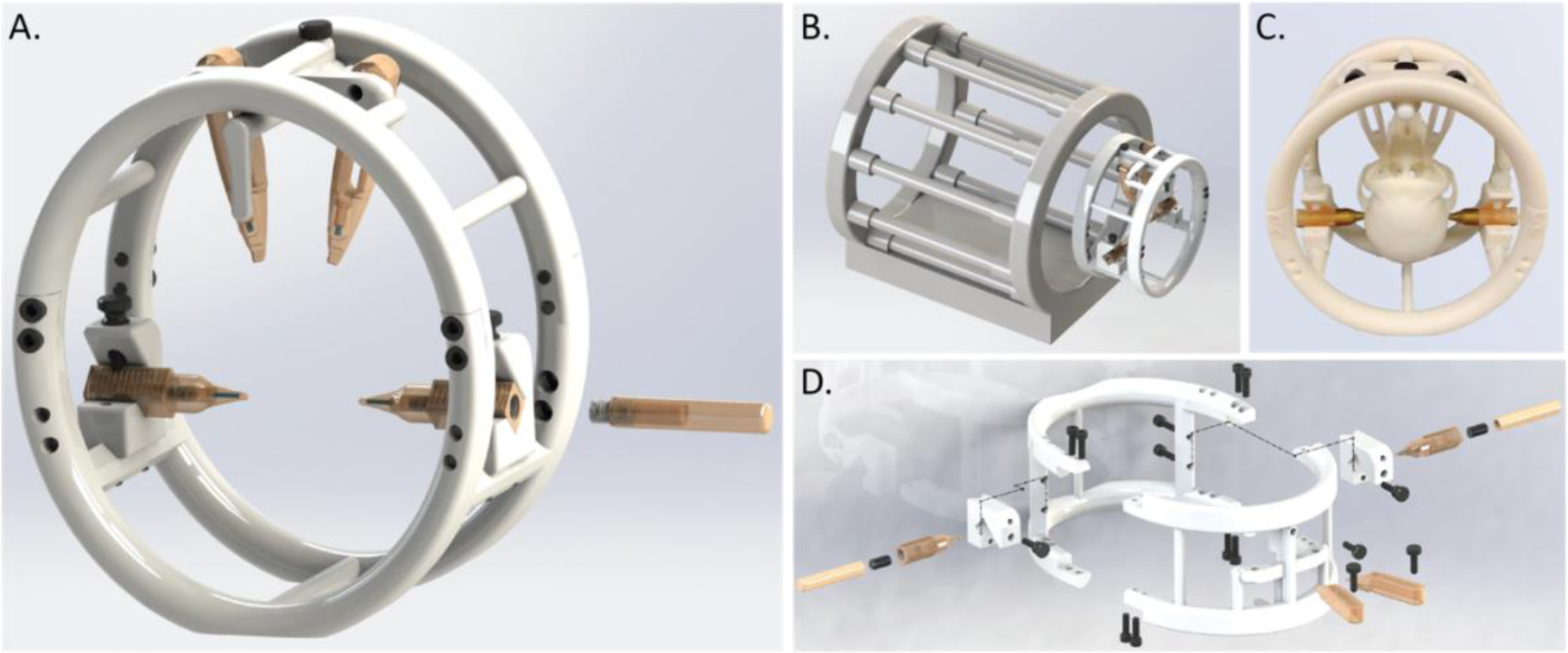
Frame design. **A)** Three-dimensional rendering of assembled frame. Right ear bar shows a removable ear bar extension that can be attached as the animal is fitted. **B)** Frame next to a rendering of a standard “bird-cage” style MRI coil. **C)** A photograph of the frame, with a 3D-printed macaque skull fitted in supine position inside the frame. **D)** A rendering of frame components, expanded for ease of view.

**Figure 2.**
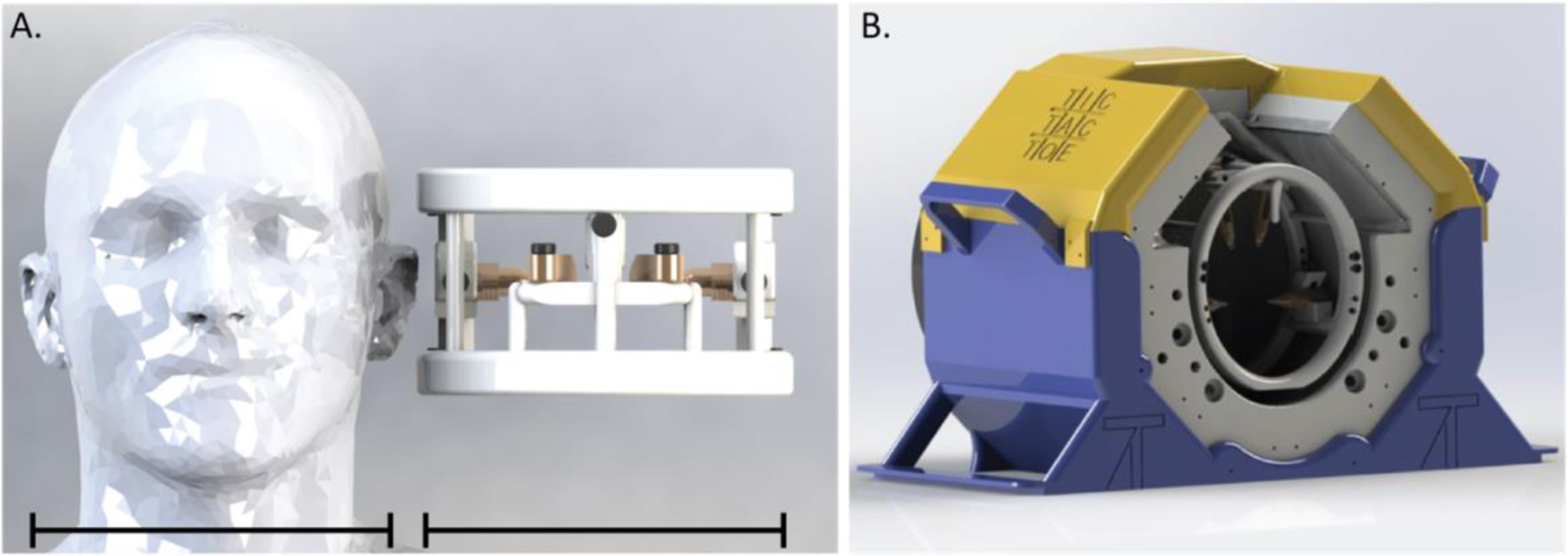
Frame size and fit. **A)** The frame next to a human head model (addapted from model published in Christ et al., 2009). **B)** Frame fitted into 2^nd^ generation Tic-tac-toe radiofrequency coil for the Siemens 7T scanner (Sajewski et al., 2023; Santini et al., 2021).

For the experiments described herein, the majority of the frame parts were 3D printed using a stereolithography printer (White Resin V4; Form 3, Formlabs, Somerville, Massachusetts, USA). Formlabs White V4 resin was chosen for its rigidity, MRI compatibility (see Image Quality section to follow) and ease of tapping. For the ear bars used with the smaller NHP species (*Cebus apella*), we opted to machine the ear bar tips to allow for placement in smaller ear canals while also minimizing wall thickness and the distance between the fiducial marker and ear bar tip. Duratron U1000 (Mitsubishi Chemical Advanced Materials) was chosen for rigidity and MRI compatibility and milled with a desktop mill (Roland MDX-50, Roland DGA, Irvine, California, USA) with a 1/8” square bit for roughing and a 1/16” ball-end bit for finishing. The use of a rigid resin (e.g., Formlabs Rigid 10k) achieves a similar result.

### 2.3 Image Quality Assessment

As an index of MRI-compatibility, we measured geometric distortion of an agarose-based MRI phantom as compared to a CT of that same phantom. We also compared the signal-to-noise ratio (SNR) of the geometric phantom with and without the stereotax inserted into the head coil. Specifically, we used a Bruker BioSpin MRI GmbH phantom (CuSO_4_ x 2H_2_O 1g/L, Agar/Agar 10g/L, model no. 1P T11170, Ettlingen, Germany) with cylindrical shape of size comparable to a monkey’s head (i.e., radius = 35 mm and height = 190 mm, **Fig. 3A**) and internal geometry of size comparable to a monkey’s brain (46×46.5×77.5 mm, **Fig. 3A**). For both assessments, MRI images were acquired in a 7T MRI (Magnetom, Siemens Healthcare, Erlangen, Germany) with a gradient-echo sequence with the following parameters: TE/TR=8.16/40 ms, resolution 400 μm isotropic, matrix size=176×384×384, flip angle=15°, acceleration factor (GRAPPA)=2. SNR maps were calculated by dividing the signal in the image by the standard deviation of the background noise. CT images (200 μm isotropic, FOV: 79.6×199 mm) of the same phantom were acquired using a Low Dose 1 mm aluminum filter and a “step and shoot” method (0.6-degree gantry step) and reconstructed using the filtered back projection algorithm in a small animal CT scanner (Si78; Bruker BioSpin GmbH, Ettlingen, Germany) equipped with the software package Paravision-360 (version 3.2; Bruker BioSpin Corp, Billerica, MA). Geometric distortions were assessed by skeletonizing (i.e., binarizing the intensity to include only the internal structure) the CT which was manually registered to the MRI, then comparing the rectilinear internal structure of the phantom to the MRI SNR maps with and without the stereotax in the coil. Importantly, the phantom was positioned in the coil with the same orientation for both scans.

**Figure 3.**
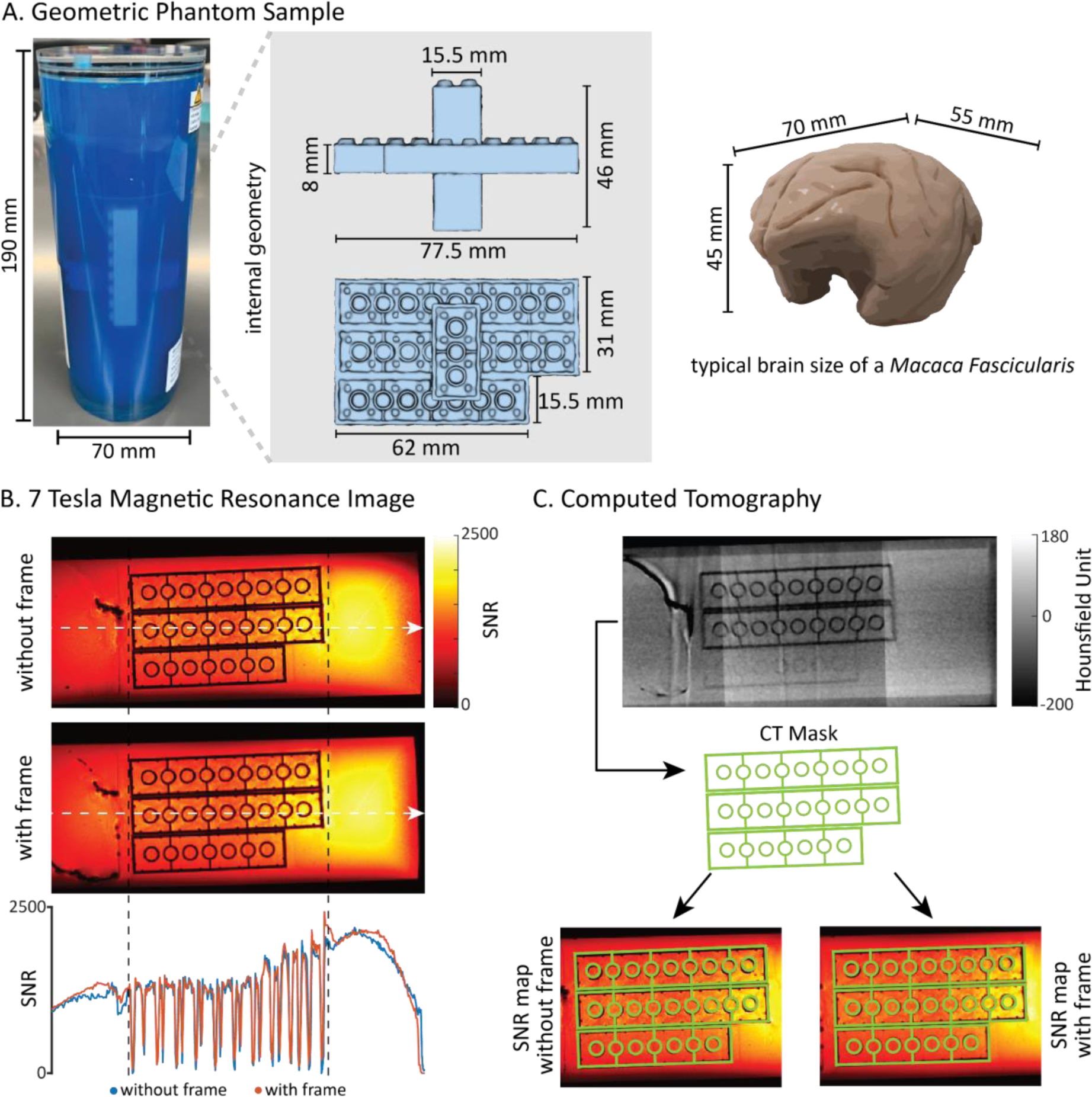
Assessment of signal quality and distortion using a phantom sample. **A)** A photograph of the MRI phantom used, with the linear internal geometry shown and overall dimensions comparable to a typical macaque brain. **B)** Signal-to-noise-ratio (SNR) maps acquired at 7 Tesla MRI (with the same hardware as used as the monkey scans) with and without stereotaxic frame. *Bottom:* overlaid slice-wise SNR with and without the frame inserted in the coil. **C)** A CT of the phantom geometry overlaid (top) on the MRI SNR maps (bottom), demonstrating minimal geometric distortion when comparing with and without the frame in the coil.

### 2.4 In vivo MRI Settings

In vivo T1-weighted MRIs were acquired for presurgical planning at the University of Pittsburgh on a human, whole body 7T MRI scanner (400 μm isotropic, TR/TE: 6000/3.7 ms, FOV: 148×148×96 mm) and using the 2^nd^ generation of the Tic-Tac-Toe radiofrequency (RF) system, with 60 transmit channels and 32 receive channels. This RF system family provides images with excellent homogeneity, coverage, and signal to noise ratio (Sajewski et al., 2023; Santini et al., 2021). Animals (MK-EF, MK-SC, MK-SZ, MK-OP, MK-HS, MK-JC, MK-JN, MK-BB) were anesthetized and intubated (induced with Ketamine, 10 mg/kg, maintained with isoflurane, 2%) with all vital signs continuously monitored. The head of the monkey was placed in the stereotactic frame (**Fig. 1A**) by securing the ear bars, eye bars, and the palate bar. A vitamin E capsule was attached on one side of the monkey’s head for distinguishing left and right in the MR image. The frame was rigidly attached to the head coil using 3D-printed brackets and nylon screws, but the frame features a flat bottom that allows for simple placement of the frame in any head coil – centering the frame allows for optimization of SNR profiles, especially with custom array applications.

An in vivo 2DSWI scan was also acquired for MK-EF in the same scanner and transmission coil, with the same animal preparations. Using a TR/TE of 1650/14 ms, flip angle of 60 degrees, and an FOV of 128×128×79.5 mm, we acquired a scan with 154 μm in-plane resolution and 1.5 mm thickness.

### 2.5 In vivo CT Settings

To optimize contrast for bone and the plastic apparatus, a CT scan (250 μm isotropic) was acquired in an Epica Vimago GT30 animal scanner for six monkeys (MK-SC, MK-SZ, MK-OP, MK-HS, MK-JC, MK-JN). Placement of the animal within the stereotaxic frame was the same as for the MRI, in the supine position. The CT and MRI images were then rigidly aligned by aligning the fiducial markers from the MRI with the CT (i.e., the ear bar hyperintensity from the MRI was aligned within the channel as apparent on the CT, same with the eye bar positions). These easily aligned images provided additional structural information for presurgical planning of deep brain targeting based on bone fiducials (as described in section *2.7*).

Detailed methods describing in-vivo MRI and CT acquisitions using the stereotaxic frame can be found at protocols.io (dx.doi.org/10.17504/protocols.io.q26g7p7rkgwz/v1).

#### 2.6.1 Utilizing the Frame for MRI-guided Real Time Injection

We tested the adaptability of the frame by performing an adeno-associated virus (AAV) injection in the scanner using real-time updated MR images to localize the injection site in MK-EF. Specifically, the frame allowed for quick identification of coordinates of a deep target site with reference to the fiducial markers while also allowing for the removal of the animal from the head coil to insert the virus injection cannula, then placement back in the coil in the same position in the scanner. The injection procedure was based on previously described methods (Yazdan-Shahmorad et al., 2018). Briefly, baseline T1-weighted scans were performed to obtain structural information (**Fig. 6B *top left***). This was followed by a second T1-weighted scan with a “dummy” probe filled with a gadolinium solution (100:1 sterile saline:gadolinium; Hospira, SKU: 63323-186-01, Lake Forest, IL, USA; Gadavist^TM^, Bayer Healthcare Pharmaceuticals, Leverkusen, Germany) placed in the chamber grid based on the coordinates (distance from ear and eye bars in a stereotaxic plane, **Fig. 6A**) determined from the baseline scan. The injection site was determined using the location of the probe within a custom recording chamber grid relative to the distance from the ear and eye bars in a stereotaxic plane. Once the location was determined, a reflux-resistant injection cannula was lowered into position and 10 μl of AAV (titer: 1.87×10^13^ GC/ml, AAVrg-Syn-ChR2(H134R)-GFP, Addgene, 58880-AAVrg) doped with gadolinium (250:1, virus:gadolinium) was injected using the convection enhanced delivery (CED) method (Bankiewicz et al., 2000). A final 2DSWI scan was run to visualize the placement of the viral injection (**Fig. 6B *top right***). The animal was then recovered and transferred back to its home cage. A detailed video protocol for CED MR-guided injection can be found on app.jove.com (doi.org/10.3791/59232-v).

#### 2.6.2 Tissue Processing and Histology

After viral injection, the animal survived for 7 months allowing sufficient time for transduction and subsequent fluorescent protein expression. The animal was euthanized and perfused transcardially with phosphate-buffered Saline (PBS, pH 7.2-7.4), followed by a series of fixatives (4% paraformaldehyde, 4% paraformaldehyde with 10% sucrose). Following cryoprotection, the brain was blocked in the coronal plane, frozen, and cut on a Leica CM1950 cryostat (RRID:SCR_018061) in 50 µm sections. Serial free-floating brain sections were selected to provide a full representation of the spread of viral injections as well as identifying injection sites. Sections selected for GFP were rinsed in Phosphate-Buffered Saline (PBS, pH 7.2-7.4). They were then moved into a 0.3% H_2_O_2_ buffer for 30 minutes, followed by rinses in PBS before incubating in a Normal Goat Serum blocking solution for one hour. The tissue was transferred to primary antibody solution (Anti-GFP 1:10000; Thermo Fisher, Molecular Probes Cat# A11122, RRID:AB_221569) for overnight incubation at 4°C. The sections were again rinsed in PBS and incubated in secondary biotinylated goat anti-rabbit IgG antibody (Vector, Vectastain ABC, Peroxidase Kit, PK-4001, 1:200) for 30 minutes. Sections were rinsed in PBS and placed into ABC (Vector, Vectastain ABC, Peroxidase Kit, PK-4001) for one hour before being rinsed in PBS. Tissue was placed in DAB substrate (3,3’-diaminobenzidine, Vector, SK-4100) for up to 6 minutes. Following a final rinse, sections were mounted and cover slipped with cytoseal before imaging (Nikon Eclipse Ti2-E microscope) and analysis (ImageJ, RRID:SCR_003071, Schneider et al., 2012). The step by step histology protocol used here can be found at protocols.io (dx.doi.org/10.17504/protocols.io.eq2lyjy4mlx9/v1).

#### 2.7.1 Cross-modality Frame Alignment for Surgical Robot Electrode Insertion

The stereotactic frame was utilized for multimodal presurgical planning for the implantation of stimulating electrodes within a deep brain structure (internal capsule) in monkeys (MK-SC, MK-SZ, MK-OP, MK-HZ, MK-JC). For this experiment, it was critical that the MRI (brain) and CT (skull) were accurately co-registered since electrode implantation trajectories are calculated based on the skull position while the target of the electrode is the brain. To do this, we manually aligned the ear-bar stoppers from the CT with the (mechanically concentric) ear bar tips from the MRI (aligned in axial and coronal planes), in the ROSA ONE^®^ Brain Robot Assistance Platform software (Zimmer Biomet, Warsaw, IN, USA). We then further confirmed the alignment by visually checking the conformity between the brain outline and the internal surface of the skull. After co-registration, we planned the trajectory of the deep brain electrode to target the hand area of the corticospinal tract (CST) within the internal capsule (IC) using the ROSA software. We identified the anterior and posterior commissures (AC-PC) and rotated the fused images to align it to the x-axis of the mid sagittal plane. The posterior commissure was set as the zero point, and coordinates (L/R, Ant./Post., Sup./Inf.) of the target in the IC were obtained respectively.

Once the robot was registered to the animal’s head, the IC electrodes were then implanted and stimulated continuously at 1 Hz or 2 Hz. The threshold of the stimulation was determined by observing a movement in the hand for each monkey, with stimulations at and above threshold amplitudes on the range of 0.8-4.8 mA (for detailed parameters used, see **Supplementary Table 1**). Electromyograms (EMG) were recorded and analyzed in MATLAB (version R2020b, RRID:SCR_001622) following a procedure as previously described (Ho et al., 2022). The EMG data and code used here can be found at zenodo.org (doi.org/10.5281/zenodo.10463426).

#### 2.7.2 Assessment of Accuracy using High-resolution Postmortem MRI

To verify that the electrode placement was correct based on cross-modality (MRI-CT) alignment of the stereotactic coordinates, we acquired ex vivo MRI in all monkeys that underwent surgical robot electrode insertion (section *2.7*) to identify stimulation location at ultra-high resolution. Following completion of the intraoperative electrophysiology, we performed a thermal lesion at the IC electrode (80° C for 1 min) to determine actual implantation position. After the surgery, the animals were euthanized and perfused with 1x phosphate buffered saline (PBS), followed by 4% paraformaldehyde (PFA). The brain of the animals was extracted and placed in 4% PFA for another 24 hours. We collected T2 (125 μm isotropic, TR/TE:1500/60ms, FOV: 52×80×56 mm) and T2* weighted images (80 μm isotropic, TR/TE: 100/16 ms, FOV: 55×70×45 mm) of the postmortem brain in a 9.4T/31 cm horizontal-bore Bruker AV3 HD animal scanner.

To calculate the target implant error, we first aligned the AC-PC in the postmortem MRIs to the x-axis of the mid sagittal plane, and set PC as the zero point, just like for the in-vivo MRI, to put them in the same coordinates. We then identified the tip of the electrode at the center of the lesion within the IC and obtained its pixel coordinates. We then converted the coordinates to metric distance (mm) based on the respective pixel dimensions. The target implant error (IE) was calculated as the Euclidean distance 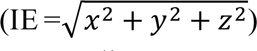 between the coordinates of the electrode tip and the ROSA robot planned target coordinate.

### 2.9 Multimodal compatibility

As proof-of-concept that the frame can be used for positron emission tomogoraphy (PET), the ear and eye bars were filled with a small dose (0.05 MBq) of [^18^F] fluorodeoxyglucose (FDG) and imaged in a small preclinical PET/CT scanner (Si78; Bruker BioSpin GmbH, Ettlingen, Germany) equipped with the software package Paravision-360 (version 3.2; Bruker BioSpin Corp, Billerica, MA, RRID: SCR_001964). Similarly, the eye bars were filled with a small FDG dose (1.54 MBq) to allow for visualization in the PET – the doses here were only for proof of concept. Practically, even smaller doses of a radioligand would be sufficient if exposure or other technical concerns with the number of detected events emerge. Using a Low Dose 1 mm aluminum filter, a 200 μm isotropic (FOV: 79.6×81.1 mm) CT was acquired using a “step and shoot” method (0.6-degree gantry step) and reconstructed using the filtered back projection algorithm. For the eye bars, a dose of 1.54 MBq was administered, while for the ear bars, a dose of 0.05 MBq was administered.

## 3. Results & Discussion

### 3.1 Frame Design

The resulting frame design is comparable to the size of a human head, with all fixation parts (eye bars, ear bars, pallet bar) fitting within the perimeter of the frame (**Fig. 2A**). This allows the frame to fit in most clinical MRI scanners either with a commercial or custom designed radio frequency coil (see **Fig. 2B** for the custom high-sensitivity conformal coil used here, Krishnamurthy et al., 2019; Sajewski et al., 2023; Santini et al., 2021, 2018). The CAD designs have been made freely and publicly available with the hope of prompting open collaboration and supporting the 3R’s in NHP research (Russell and Burch, 1959). Each component is provided as a stereolithography (STL) file as well as a Standard for The Exchange of Product (STEP) file, ready for 3D-printing or machining (doi.org/10.5281/zenodo.10456840). Vendor-agnostic initial-graphics-exchange-specification (IGES) files are also included and can be imported into freely available CAD software for modification, if desired. Engineering drawings and a bill of materials are provided to specify tapped holes and required MRI-compatible hardware (e.g., nylon screws, **Supplementary Fig. 1 and 2)**. A variety of sized ear, eye, and pallet bar designs are included to accommodate a variety of NHP species and head sizes – for example capuchin monkeys require a much longer eye and pallet bars than do macaques. Importantly, the frame has a modular design that allows printing of each piece on small format desktop 3D printers (such as Form 3/3B+). A full set of frame components (**Supplementary Fig. 1**, items 1-8) can be printed within a few days, and for a total cost of less than $75 USD in 2023 (time and cost details for each component reported in **Supplementary Table 2**).

In order to assess MRI compatibility of the frame, we compared the MRI signal-to-noise ratio (SNR) of an MRI phantom with known internal geometry (**Fig. 3A**) with and without the frame inserted into the coil (**Fig. 3B**). This geometry was then compared to a CT image of the same phantom (**Fig. 3C**). Minimal variation (< 3 %) was observed in the SNR (average of 369 verses 379) when the frame was used, likely due to small variation in the positioning of the phantom inside the coil. Importantly, there was also minimal susceptibility distortion induced by the frame (**Fig. 3C**).

Finally, we highlighted the ability to reliably scan animals of different sizes and species across different modalities. We demonstrate that the frame can be used on a variety of NHP species (*Macaca mulatta*, *Macaca fascicularis*, and *Cebus apella*) whose weights range from 2.93 kg (**Fig. 4A**) to over 10kg (**Fig. 4C**).

**Figure 4.**
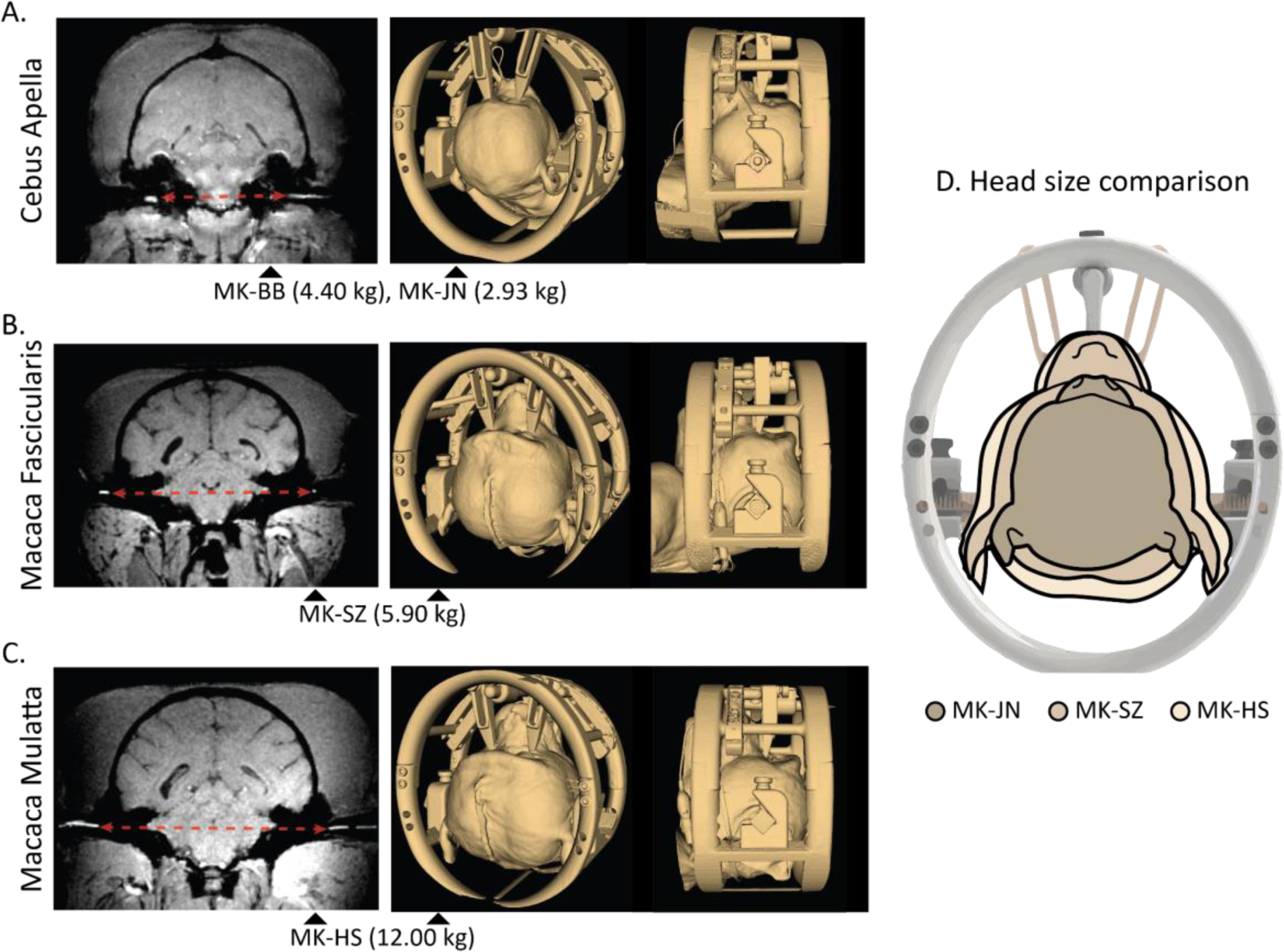
Examples of different species and weights scanned with the frame. 7T MRI (left) showing CuSO_4_ ear bar fiducials within the external auditory meatus (alignment shown with red arrow), as well as CT segmented top and side renderings of monkey inside frame (right) for **A)** MK-BB & MK-JN, Cebus apella, 4.40 kg & 2.93 kg; **B**) MK-SZ, Macaca fascicularis, 5.90 kg; **C)** MK-HS, Macaca mulatta, 12.00 kg. **D)** Comparison of head size of the 3 monkeys inside frame.

### 3.2 Cross-modality adaptability

The ear bars tip can be filled with a copper sulfate solution (CuSO_4_, **Fig. 5A**), which allowed its visualization in the T1-(and T2-weighted) MRI contrasts in the MRI (**Fig. 5C**). Filling the fiducial tips with [^18^F] fluorodeoxyglucose (FDG) also allowed visualization of the fiducial chamber in PET scans, which can be acquired together with CT scans without additional registration (**Fig. 5B**). Additionally, the nylon thread for ear-bar extension bars (concentric with the fiducial channels) has a radio-density similar to bone and can be easily visualized in the CT bone window of a monkey (**Fig. 5D**). These features together reduced the degrees of freedom in transformation needed during co-registration from 6 (3x translation & 3x rotation) to 4 (3x translation & coronal plane rotation), ensuring both registration efficiency and accuracy (**Fig. 5D**). Co-registration results were approved by an experienced neurosurgeon.

**Figure 5.**
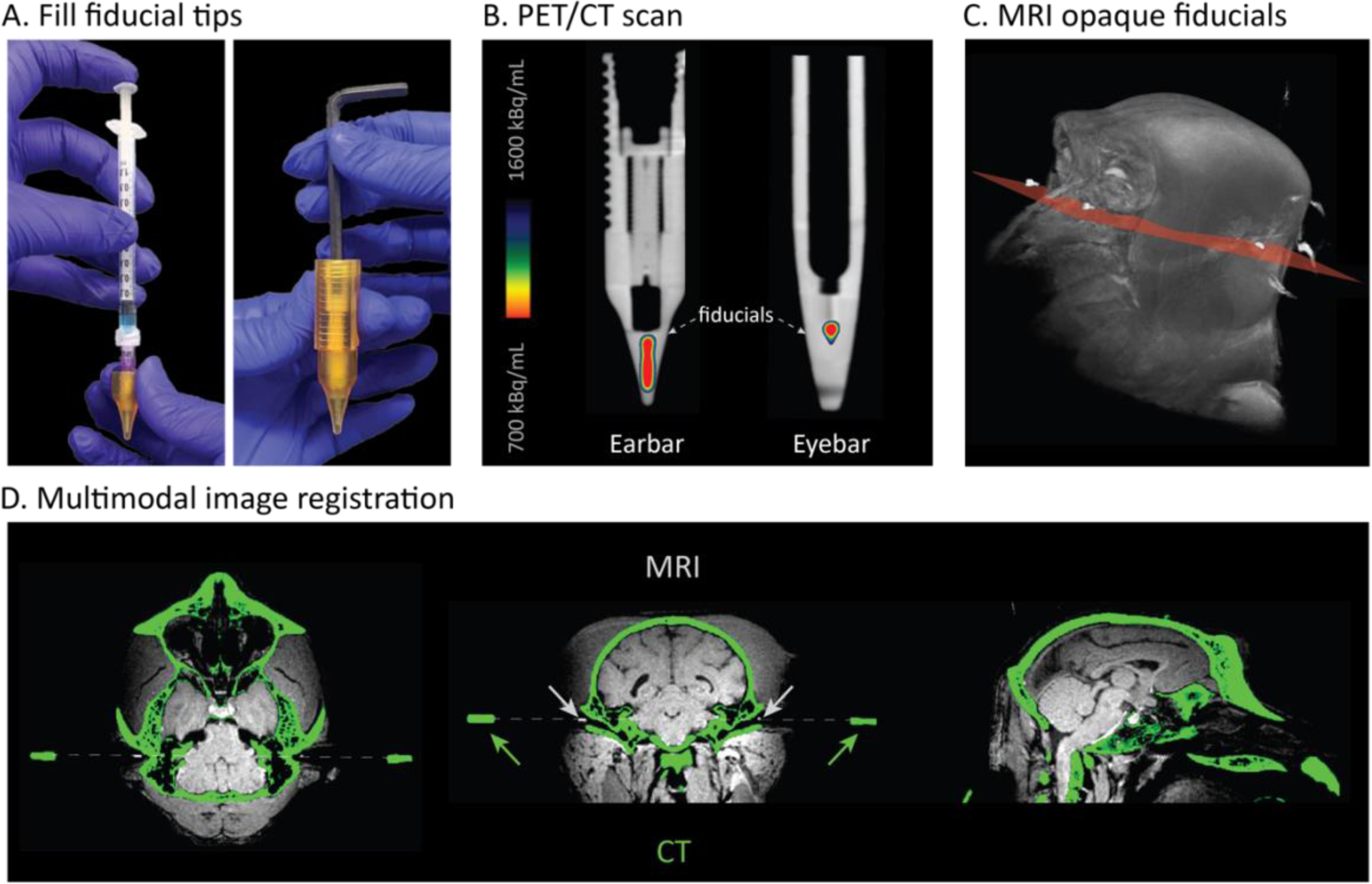
Cross-modality alignment. **A)** Demonstration of filling an ear bar fiducial tip with a syringe and sealing the chamber with a MRI-compatible nylon screw. **B)** A FDG-PET scan overlaid on a CT scan, showing the FDG filled fiducial chambers. **C)** An MRI reconstruction of the monkey head with CuSO_4_ fiducials in the ear bars and eye bars visible and aligned in the stereotaxic plane. **D)** A CT-scan (green) overlaid with an MRI (grey). The Copper Sulfate (CuSO_4_) filled ear bars are highlighted to show ability to align across multiple imaging formats.

### 3.3 MR-guided injection

We demonstrate the experimental applicability of our low-cost 3D-printable frame by first assessing the accuracy of AAV injection for neuronal tracing. For this, an animal was fitted within the frame and T1-weighted MRI image was acquired to determine the anatomical target (caudate nucleus) with reference to the fiducial markers. Based on that location, a probe filled with 2% Gadolinium was then inserted in the brain and a second scan was acquired to confirm injection location (caudate nucleus). Once the location was confirmed, the AAV doped with Gadoteridol was infused and a final scan was acquired to visualize the placement of the viral injection (**Fig. 6**). Histology showed a clear injection site that overlapped with the visible site in the MRI within the caudate nucleus (Cd, **Fig. 6B *mid left and right***). Systematic serial labeling of sections also shows an injection site of roughly 3 mm^3^. Despite some injection tract reflux into the IC, systematic immunohistochemical (IHC) staining for GFP revealed successful viral transport, primarily to cortical layer 5 neurons located throughout the prefrontal cortex including Brodmann Area (BA) 8, 6, and 46 primarily limited to layer 5 (**Fig. 6B *bottom***). In summary, the IHC labeling demonstrate the ability to target subcortical structures and reliably maintain a fiducial coordinate system, allowing for deliver viral tracers or agents while the monkey is fixed in the stereotactic apparatus.

**Figure 6.**
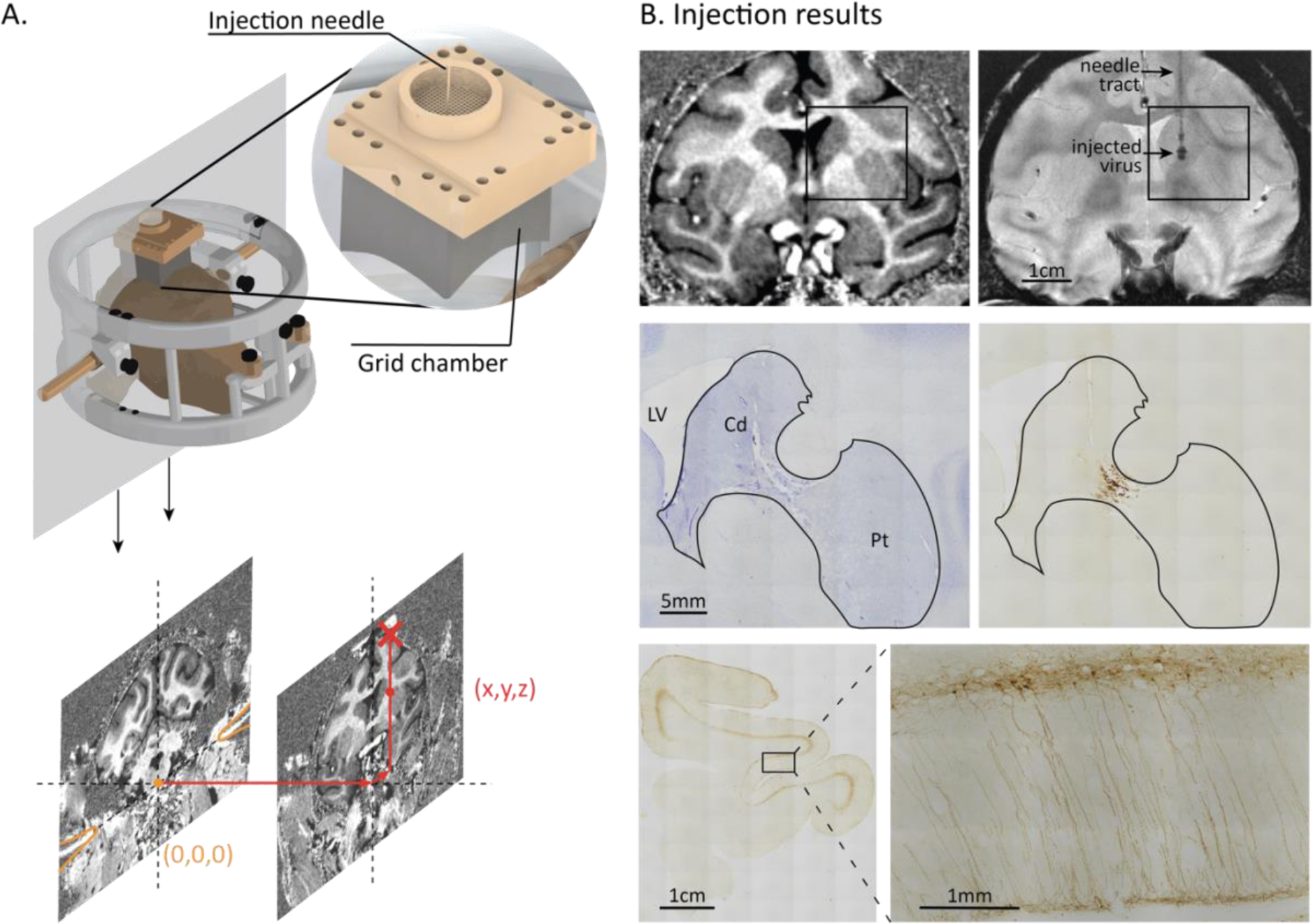
Viral expression of rAAV-GFP in striatum and cortex. **A) *Top:*** Rendering of MK-EF inside the stereotaxic frame with an enlargement of the grid chamber and the cannula used for the rAAV injections. ***Bottom:*** Coronal T1-weighted MRI slice aligned to the ear bar (yellow) shows stereotaxic origin (0,0,0), and slice aligned with injection target (red) showing respective stereotaxic target coordinates (x,y,z), as well as chamber location (red cross). **B) *Top left:*** T1-weighted MRI prior to injection. ***Top right:*** *2DSWI* MRI showing location of viral injection within the caudate nucleus. ***Mid left:*** Nissl-stained section showing injection site. ***Mid right:*** Adjacent section to that shown in *Mid left* stained for GFP highlighting spread of the infection. ***Bottom left:*** Example of cortical labeling in the ipsilateral hemisphere to the injection. ***Bottom right:*** Area within the box in *Bottom left* highlighting the layer specificity of the viral projections.

### 3.4 Multimodal alignment for deep brain electrode implantation and validation

The stereotactic frame was utilized for multimodal presurgical planning for the implantation of stimulating electrodes within a deep brain structure (internal capsule) in five monkeys. For this experiment, it was critical that the MRI (brain) and CT (skull) were accurately co-registered, with the electrode implantation trajectories calculated based on the skull position while the target of the electrode is the brain. To do this, we manually aligned the ear bar position from the CT with the (mechanically concentric) ear bar tips from the MRI (aligned in axial and coronal planes), in the ROSA ONE^®^ Brain Robot Assistance Platform software (Zimmer Biomet, Warsaw, IN, USA).

Following completion of registration of the surgical robot to each bone fiducial on the animal’s skull, based on 3D marker coordinates defined with the CT images, the robot reports a registration error. The average target registration error (TRE, reported by the ROSA ONE^®^ Brain software) was 0.316 mm for the five monkeys (MK-SC: 0.39 mm, MK-SZ: 0.32 mm, MK-OP: 0.25 mm, MK-HS: 0.29 mm, MK-JC: 0.33 mm. **Fig. 7A**), which is comparable to registration error in human surgeries (González-Martínez et al., 2016; Xu et al., 2018). While stimulating the internal capsule, we recorded motor evoked potentials in hand, arm, and face muscles. IC stimulation at threshold amplitude (MK-SC: 4.5 mA, MK-OP: 1 mA, MK-HS: 1 mA, MK-JC: 1.8 mA) elicited responses in the hand muscles (flexor digiti minimi and/or abductor pollicis brevis) but no responses in the lip or biceps, indicating the accurate placement of the IC probe within the hand area of the IC (**Fig. 7B**). At suprathreshold amplitude (MK-SC: 4.8 mA, MK-OP: 1.5 mA, MK-HS: 2 mA, MK-JC: 2 mA), we observed a larger response in the hand muscles and still no responses in the lip or biceps.

**Figure 7.**
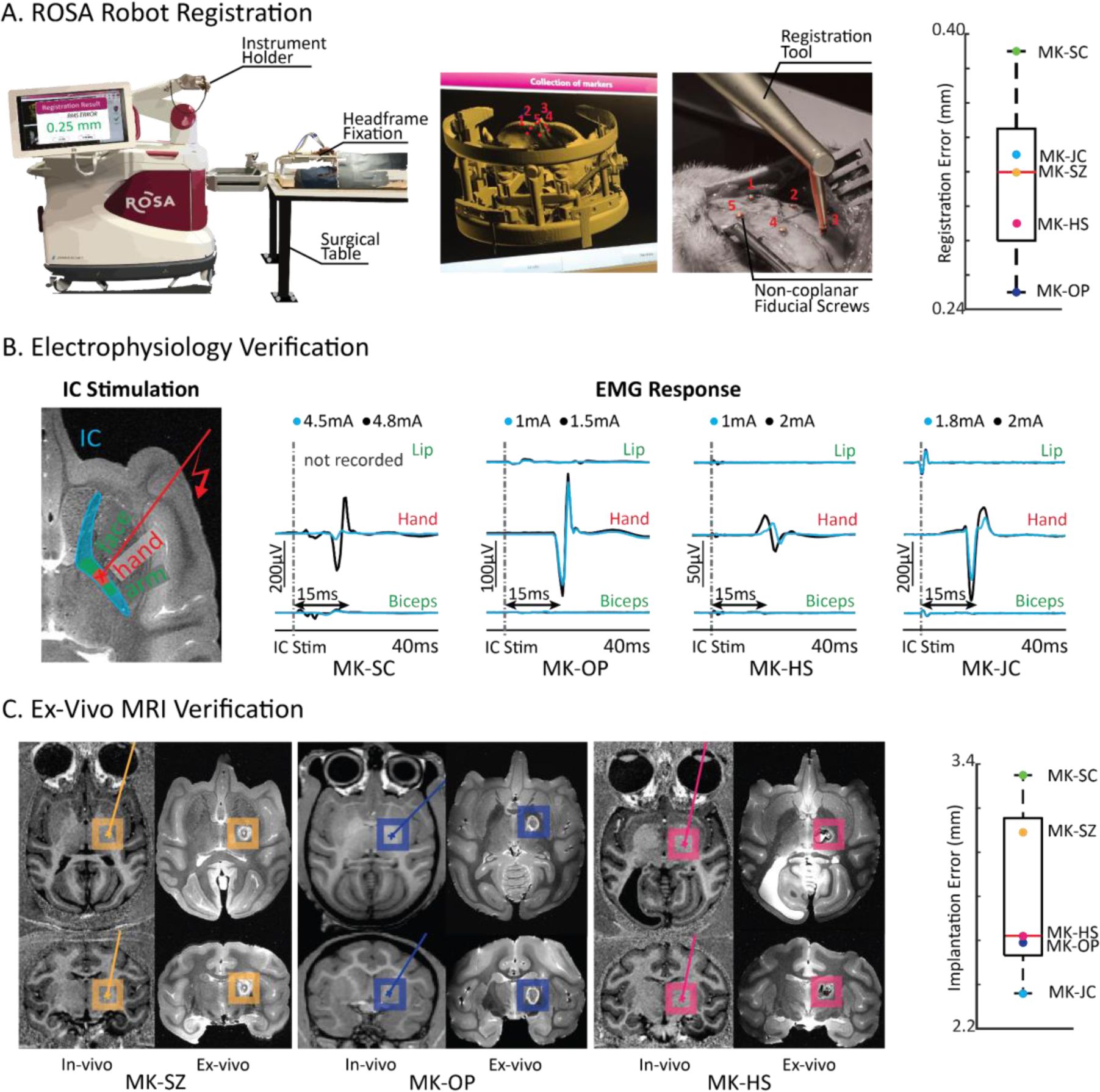
Robot assisted deep brain electrode implant validation. **A)** Demonstration of ROSA robot registration between merged MRI/CT images and titanium skull fiducial screws, with target registration errors (TRE) shown on the right for each monkey. **B)** Electrical stimulation through hand IC targeting electrode (left). Stimulation evoked EMG responses (n=4 monkeys) localize to hand muscles, and not face (lip) or arm (biceps) muscles. **C)** Examples of MRIs comparing planned targets (in-vivo) and implant lesions (ex-vivo). Implantation error between the two are shown for each monkey.

For ex-vivo MRI verification of the implanted IC electrode, we calculated Euclidean implantation errors between the planned IC target and the tip of the lesion in the post-mortem T2-weighted MRI. The mean implantation error was 2.74 mm for 5 monkeys (MK-SC: 3.43 mm, MK-SZ: 2.75 mm, MK-OP: 2.54 mm, MK-HS: 2.62 mm, MK-JC: 2.36 mm, **Fig. 7C**) in line with human surgeries as well as reported high precision monkey implants (González-Martínez et al., 2016; Zhu et al., 2019).

Overall, these results confirm an accurate implantation of deep brain electrodes that was possible thanks to a precise co-registration of CT and MRI images that was made possible by the use of our customized MRI-compatible stereotaxic frame that readily fit in clinical MRI and CT scanners.

## 4. Conclusions

Here, we developed a compact MRI-compatible stereotaxis suitable for a variety of NHP species (*Macaca mulatta*, *Macaca fascicularis*, and *Cebus apella* shown here) that allows for multimodal alignment through technique-specific fiducial markers. The design is compatible with most clinical neuroimaging (MRI, PET, CT) systems, and that can be easily manufactured using a standard stereolithography 3D printer. The frame is intended to be used across modalities, with liquid-tight and refillable fiducial reservoirs for modality-specific contrast enhancement. The lightweight build allows for easy maneuverability when placing the animal in the scanner of choice while maintaining the proper planer alignment. The fiducial markers provide key points of reference from which reliable coordinates can be determined and accurate coregistration across imaging modalities is favored. In this paper, we demonstrate practical utilization of the frame enabling accurate real time MR-guided deep brain viral injections, as well as electrode implant trajectory planning with multimodal imaging targeting the internal capsule and validated through electrophysiology. We have also used the frame to successfully implant specific nuclei within the thalamus, with which we studied their roles in facilitating motor control after stroke through electrophysiology experiments (data presented in Ho et al., 2022). With these results we demonstrate that this deep brain surgery planning approach using our 3D printed stereotaxic frame can be applied for any other deep brain injection and electrophysiological studies.

## Data and Code Availability

Stereotaxic frame model files can be found at zenodo.org (doi.org/10.5281/zenodo.10456840) and github.com (https://github.com/isabelazr/Macaque-Stereotax.git). EMG data and code were uploaded to zenodo.org (doi.org/10.5281/zenodo.10463426). Detailed protocol for In-vivo MRI and CT can be found at protocols.io (dx.doi.org/10.17504/protocols.io.q26g7p7rkgwz/v1). A video protocol for CED MR-guided injection can be found at jove.com (doi.org/10.3791/59232- v). The histology protocols used in this paper can also be found at protocols.io (dx.doi.org/10.17504/protocols.io.eq2lyjy4mlx9/v1).

## Author Contributions

D.J.S. conceived the study. D.J.S. and E.P. secured funding. D.J.S. designed the stereotaxic frame. D.J.S., A.A., A.C.B, W.R.S. performed the MR guided injections and histological processing. L.L., J.H., E.P. designed and performed the robotic deep brain implantation and validation experiments. L.L., A.A., J.H., I.Z.R, T.S. created the figures. L.L., I.Z.R, D.J.S., E.P. wrote the paper, and all authors contributed to its editing. All authors approved the final version of the manuscript and agreed to be accountable for all aspects of the work in ensuring that questions related to the accuracy or integrity of any part of the work are appropriately investigated and resolved. All persons designated as authors qualify for authorship, and all those who qualify for authorship are listed.

## Acknowledgement

We wish to thank Zimmer Biomet for lending the ROSA robot to use in our surgeries (we declare no conflict of interest). We would also like to thank Tyler Simpson for his help summarizing print time and cost, as well as advice from an engineering perspective. We also wish to thank Dr. Julia Oluoch, Erica Griffith, Stacey Cashman, Jacquelyn Breter, and Baylie Leveto for their expertise in animal handling and critical input on the design to assure animal comfort and safety. Additionally, this investigation used resources that were supported by the Southwest National Primate Research Center grant P51 OD011133 from the Office of Research Infrastructure Programs, National Institutes of Health and the NIH grant U42 OD120442.

## Appendix

**Supplementary Figure 1.**
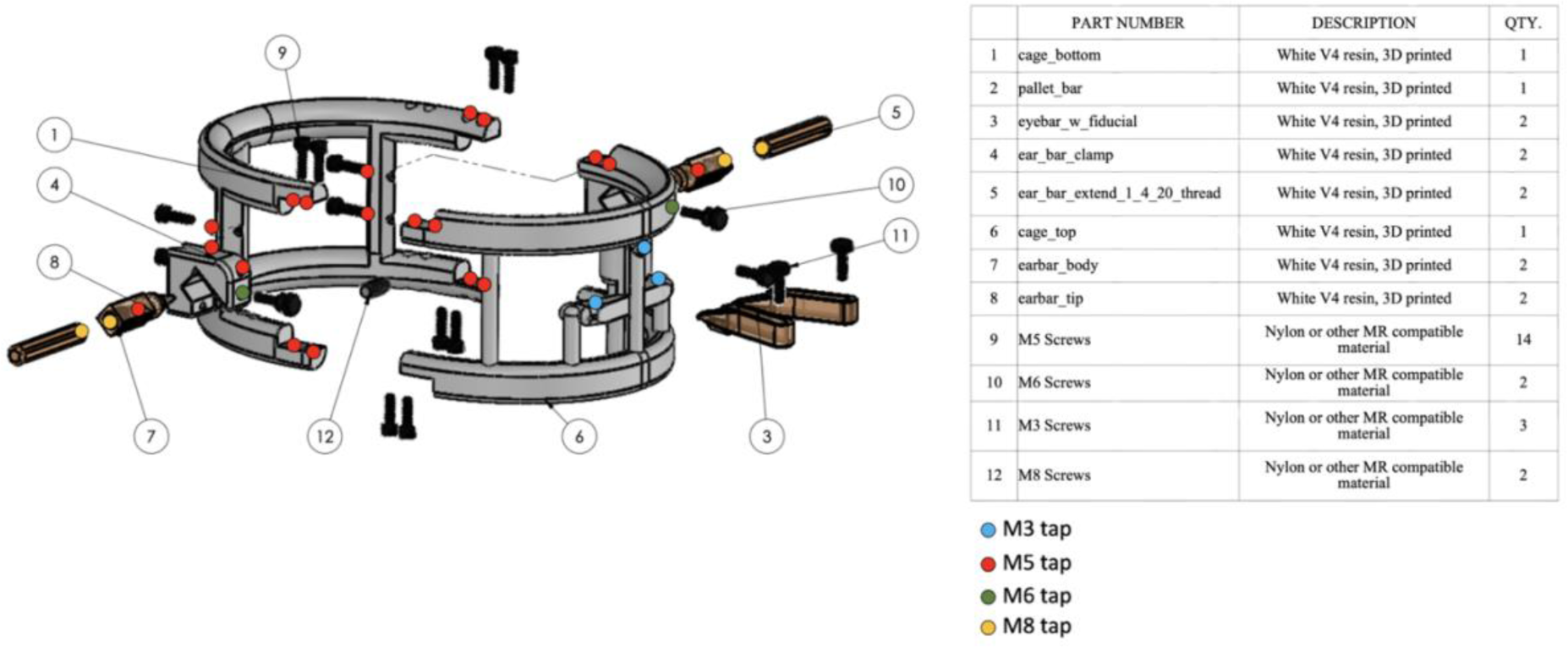
Tapping instructions and bill of materials.

**Supplementary Figure 2.**
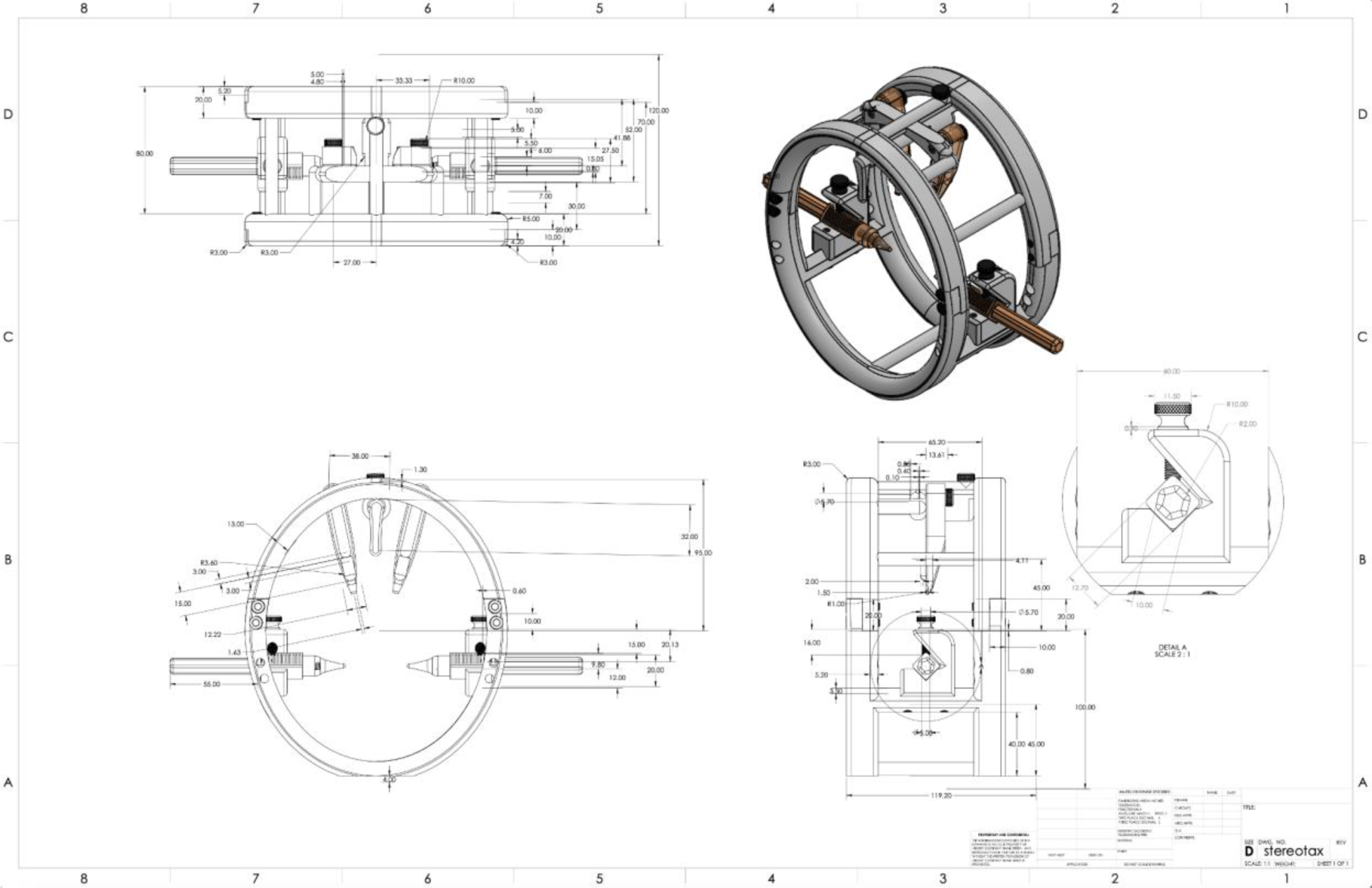
Engineering drawing.

**Supplementary Table 1.** IC Stimulation Parameters.

| Animal ID | Amplitude | Pulse Width | Frequency |
| --- | --- | --- | --- |
| MK-SC | 4.5 mA, 4.8 mA | 2 ms | 1 Hz |
| MK-SZ | 0.8 mA, 1.1 mA | 0.8 ms | 2 Hz |
| MK-OP | 1.0 mA, 1.5 mA | 3 ms | 2 Hz |
| MK-HS | 1.0 mA, 2.0 mA | 1 ms | 2 Hz |
| MK-JC | 1.8 mA, 2.0 mA | 1 ms | 1 Hz |

**Supplementary Table 2.** Modular 3D printing time and cost estimate for Form 3B+. Min means just the part, no additional support material considered. Avg means just importing the STL part into PreForm without additional adjustments, including support material with density of 0.6 and touchpoint between 0.3mm-0.5mm. Only cells bolded and in italics are used to estimate total costs.

| Part | Print time<br>(Min hr) | Print time<br>(Avg hr) | Material volume<br>(Min mL) | Material volume<br>(Avg mL) |  |  |
| --- | --- | --- | --- | --- | --- | --- |
| Cage_bottom | 9.5 | **13.75** | 142 | **186** | *Avg. time* | **41 hr** |
| Cage_top | 9.5 | **14** | 145 | **191** | *Avg. cost* | **65.649 dollars** |
| Ear_bar_clamp (for each) | 1.5 | **2.5** | 17 | **20** | *Avg. vol.* | **256.6 mL** |
| Ear_bar_extend (for each) | 0.75 | **2.75** | 4.4 | **6.3** |  |  |
| Pallet_bar | 0.75 | **2.75** | 6.6 | **11** |  |  |
| Pallet_bar_large | 0.75 | 2.75 | 6.6 | 11 |  |  |
| Pallet_bar_small | 0.75 | 2.75 | 6.5 | 11 |  |  |
| Pallet_bar_xl | 0.75 | 2.75 | 6.7 | 11 |  |  |
| Pallet_bar_xs | 0.75 | 2.75 | 6.5 | 11 |  |  |

**Material: Rigid 10k** \$299/L
| Part (for each) | Print time<br>(Min hr) | Print time<br>(Avg hr) | Material volume<br>(Min mL) | Material volume<br>(Avg mL) |  |  |
| --- | --- | --- | --- | --- | --- | --- |
| Earbar_body | 1 | **1.75** | 3.3 | **4.9** | *Avg. time* | **14 hr** |
| Earbar_tip_large | 1.5 | 1.75 | 1.7 | 2.6 | *Avg. cost* | **8.671 dollars** |
| Earbar_tip_small | 1.25 | 1.75 | 1.6 | 2.5 | *Avg. vol.* | **29 mL** |
| Earbar_tip_xl | 1.5 | 1.75 | 1.7 | 2.7 |  |  |
| Earbar_tip_xs | 1.25 | 1.5 | 1.5 | 2.5 |  |  |
| Earbar_tip | 1.5 | **1.75** | 1.7 | **2.6** |  |  |
| Eyebar_w_fiducial_large | 1 | 3.5 | 5 | 6.9 |  |  |
| Eyebar_w_fiducial_small | 1 | 3.5 | 5 | 7.9 |  |  |
| Eyebar_w_fiducial_xl | 1 | 3.5 | 5 | 8.1 | *Tot. time* | **55 hr** |
| Eyebar_w_fiducial_xs | 1 | 3.5 | 5 | 7.8 | *Tot. cost* | **74.32 dollars** |
| Eyebar_w_fiducial | 1 | **3.5** | 5 | **7** | *Tot. vol.* | **285.6 mL** |

## References

Alvarez-Royo, P., Clower, R.P., Zola-Morgan, S., Squire, L.R., 1991. Stereotaxic lesions of the hippocampus in monkeys: determination of surgical coordinates and analysis of lesions using magnetic resonance imaging. J Neurosci Methods 38, 223–232. 10.1016/0165-0270(91)90172-v

Bankiewicz, K.S., Eberling, J.L., Kohutnicka, M., Jagust, W., Pivirotto, P., Bringas, J., Cunningham, J., Budinger, T.F., Harvey-White, J., 2000. Convection-Enhanced Delivery of AAV Vector in Parkinsonian Monkeys; In Vivo Detection of Gene Expression and Restoration of Dopaminergic Function Using Pro-drug Approach. Experimental Neurology 164, 2–14. 10.1006/exnr.2000.7408

Bentley, J.N., Khalsa, S.S.S., Kobylarek, M., Schroeder, K.E., Chen, K., Bergin, I.L., Tat, D.M., Chestek, C.A., Patil, P.G., 2018. A simple, inexpensive method for subcortical stereotactic targeting in nonhuman primates. Journal of Neuroscience Methods 305, 89–97. 10.1016/j.jneumeth.2018.05.007

Chen, L., Li, N., Gao, L., Yang, C., Fang, W., Wang, X., Gao, G., 2015. Improved stereotactic procedure enhances the accuracy of deep brain stimulation electrode implantation in non-human primates. International Journal of Neuroscience 125, 380–389. 10.3109/00207454.2014.940524

Christ, A., Kainz, W., Hahn, E.G., Honegger, K., Zefferer, M., Neufeld, E., Rascher, W., Janka, R., Bautz, W., Chen, J., 2009. The Virtual Family—development of surface-based anatomical models of two adults and two children for dosimetric simulations. Physics in Medicine & Biology 55, N23.

Dubowitz, D.J., Scadeng, M., 2011. A frameless stereotaxic MRI technique for macaque neuroscience studies. Open Neuroimag J 5, 198–205. 10.2174/1874440001105010198

Fox, A.S., Holley, D., Klink, P.C., Arbuckle, S.A., Barnes, C.A., Diedrichsen, J., Kwok, S.C., Kyle, C., Pruszynski, J.A., Seidlitz, J., Zhou, X., Poldrack, R.A., Gorgolewski, K.J., 2021. Sharing voxelwise neuroimaging results from rhesus monkeys and other species with Neurovault. NeuroImage 225, 117518. 10.1016/j.neuroimage.2020.117518

Fredericks, J.M., Dash, K.E., Jaskot, E.M., Bennett, T.W., Lerchner, W., Dold, G., Ide, D., Cummins, A.C., Der Minassian, V.H., Turchi, J.N., Richmond, B.J., Eldridge, M.A.G., 2020. Methods for mechanical delivery of viral vectors into rhesus monkey brain. Journal of Neuroscience Methods 339, 108730. 10.1016/j.jneumeth.2020.108730

Frey, S., Comeau, R., Hynes, B., Mackey, S., Petrides, M., 2004. Frameless stereotaxy in the nonhuman primate. NeuroImage 23, 1226–1234. 10.1016/j.neuroimage.2004.07.001

Frey, S., Pandya, D.N., Chakravarty, M.M., Bailey, L., Petrides, M., Collins, D.L., 2011. An MRI based average macaque monkey stereotaxic atlas and space (MNI monkey space). NeuroImage 55, 1435–1442. 10.1016/j.neuroimage.2011.01.040

Friedman, H., Ator, N., Haigwood, N., Newsome, W., Allan, J.S., Golos, T.G., Kordower, J.H., Shade, R.E., Goldberg, M.E., Bailey, M.R., Bianchi, P., 2017. The Critical Role of Nonhuman Primates in Medical Research. Pathog Immun 2, 352–365. 10.20411/pai.v2i3.186

González-Martínez, J., Bulacio, J., Thompson, S., Gale, J., Smithason, S., Najm, I., Bingaman, W., 2016. Technique, Results, and Complications Related to Robot-Assisted Stereoelectroencephalography. Neurosurgery 78, 169–180. 10.1227/NEU.0000000000001034

Ho, J.C., Liang, L., Grigsby, E.M., Balaguer, J., Karapetyan, V., Schaeffer, D.J., Silva, A.C., Hitchens, T.K., Capogrosso, M., Gerszten, P.C., Gonzalez-Martinez, J.A., Pirondini, E., 2022. Robot Assisted Neurosurgery for High-Accuracy, Minimally-Invasive Deep Brain Electrophysiology in Monkeys, in: 2022 44th Annual International Conference of the IEEE Engineering in Medicine & Biology Society (EMBC). Presented at the 2022 44th Annual International Conference of the IEEE Engineering in Medicine & Biology Society (EMBC), pp. 3115–3118. 10.1109/EMBC48229.2022.9871520

Krishnamurthy, N., Santini, T., Wood, S., Kim, J., Zhao, T., Aizenstein, H.J., Ibrahim, T.S., 2019. Computational and experimental evaluation of the Tic-Tac-Toe RF coil for 7 Tesla MRI. PLOS ONE 14, e0209663. 10.1371/journal.pone.0209663

Messinger, A., Sirmpilatze, N., Heuer, K., Loh, K.K., Mars, R.B., Sein, J., Xu, T., Glen, D., Jung, B., Seidlitz, J., Taylor, P., Toro, R., Garza-Villarreal, E.A., Sponheim, C., Wang, X., Benn, R.A., Cagna, B., Dadarwal, R., Evrard, H.C., Garcia-Saldivar, P., Giavasis, S., Hartig, R., Lepage, C., Liu, C., Majka, P., Merchant, H., Milham, M.P., Rosa, M.G.P., Tasserie, J., Uhrig, L., Margulies, D.S., Klink, P.C., 2021. A collaborative resource platform for non-human primate neuroimaging. NeuroImage 226, 117519. 10.1016/j.neuroimage.2020.117519

Milham, M., Petkov, C., Belin, P., Ben Hamed, S., Evrard, H., Fair, D., Fox, A., Froudist-Walsh, S., Hayashi, T., Kastner, S., Klink, C., Majka, P., Mars, R., Messinger, A., Poirier, C., Schroeder, C., Shmuel, A., Silva, A.C., Vanduffel, W., Van Essen, D.C., Wang, Z., Roe, A.W., Wilke, M., Xu, T., Aarabi, M.H., Adolphs, R., Ahuja, A., Alvand, A., Amiez, C., Autio, J., Azadi, R., Baeg, E., Bai, R., Bao, P., Basso, M., Behel, A.K., Bennett, Y., Bernhardt, B., Biswal, B., Boopathy, S., Boretius, S., Borra, E., Boshra, R., Buffalo, E., Cao, L., Cavanaugh, J., Celine, A., Chavez, G., Chen, L.M., Chen, X., Cheng, L., Chouinard-Decorte, F., Clavagnier, S., Cléry, J., Colcombe, S.J., Conway, B., Cordeau, M., Coulon, O., Cui, Y., Dadarwal, R., Dahnke, R., Desrochers, T., Deying, L., Dougherty, K., Doyle, H., Drzewiecki, C.M., Duyck, M., Arachchi, W.E., Elorette, C., Essamlali, A., Evans, A., Fajardo, A., Figueroa, H., Franco, A., Freches, G., Frey, S., Friedrich, P., Fujimoto, A., Fukunaga, M., Gacoin, M., Gallardo, G., Gao, L., Gao, Y., Garside, D., Garza-Villarreal, E.A., Gaudet-Trafit, M., Gerbella, M., Giavasis, S., Glen, D., Ribeiro Gomes, A.R., Torrecilla, S.G., Gozzi, A., Gulli, R., Haber, S., Hadj-Bouziane, F., Fujimoto, S.H., Hawrylycz, M., He, Q., He, Y., Heuer, K., Hiba, B., Hoffstaedter, F., Hong, S.-J., Hori, Y., Hou, Y., Howard, A., de la Iglesia-Vaya, M., Ikeda, T., Jankovic-Rapan, L., Jaramillo, J., Jedema, H.P., Jin, H., Jiang, M., Jung, B., Kagan, I., Kahn, I., Kiar, G., Kikuchi, Y., Kilavik, B., Kimura, N., Klatzmann, U., Kwok, S.C., Lai, H.-Y., Lamberton, F., Lehman, J., Li, P., Li, Xinhui, Li, Xinjian, Liang, Z., Liston, C., Little, R., Liu, C., Liu, N., Liu, Xiaojin, Liu, Xinyu, Lu, H., Loh, K.K., Madan, C., Magrou, L., Margulies, D., Mathilda, F., Mejia, S., Meng, Y., Menon, R., Meunier, D., Mitchell, A.J., Mitchell, A., Murphy, A., Mvula, T., Ortiz-Rios, M., Ortuzar Martinez, D.E., Pagani, M., Palomero-Gallagher, N., Pareek, V., Perkins, P., Ponce, F., Postans, M., Pouget, P., Qian, M., Ramirez, J. “Bene,” Raven, E., Restrepo, I., Rima, S., Rockland, K., Rodriguez, N.Y., Roger, E., Hortelano, E.R., Rosa, M., Rossi, A., Rudebeck, P., Russ, B., Sakai, T., Saleem, K.S., Sallet, J., Sawiak, S., Schaeffer, D., Schwiedrzik, C.M., Seidlitz, J., Sein, J., Sharma, J., Shen, K., Sheng, W., Shi, N.S., Shim, W.M., Simone, L., Sirmpilatze, N., Sivan, V., Song, X., Tanenbaum, A., Tasserie, J., Taylor, P., Tian, X., Toro, R., Trambaiolli, L., Upright, N., Vezoli, J., Vickery, S., Villalon, J., Wang, X., Wang, Y., Weiss, A.R., Wilson, C., Wong, T.-Y., Woo, C.-W., Wu, B., Xiao, D., Xu, A.G., Xu, D., Xufeng, Z., Yacoub, E., Ye, N., Ying, Z., Yokoyama, C., Yu, X., Yue, S., Yuheng, L., Yumeng, X., Zaldivar, D., Zhang, S., Zhao, Y., Zuo, Z., 2022. Toward next-generation primate neuroscience: A collaboration-based strategic plan for integrative neuroimaging. Neuron 110, 16–20. 10.1016/j.neuron.2021.10.015

Miocinovic, S., Zhang, J., Xu, W., Russo, G.S., Vitek, J.L., McIntyre, C.C., 2007. Stereotactic neurosurgical planning, recording, and visualization for deep brain stimulation in non-human primates. Journal of Neuroscience Methods 162, 32–41. 10.1016/j.jneumeth.2006.12.007

Ohayon, S., Tsao, D.Y., 2012. MR-guided stereotactic navigation. Journal of Neuroscience Methods 204, 389–397. 10.1016/j.jneumeth.2011.11.031

Paiva, F.B. de, Campbell, B.A., Frizon, L.A., Martin, A., Maldonado-Naranjo, A., Machado, A.G., Baker, K.B., 2020. Feasibility and performance of a frameless stereotactic system for targeting subcortical nuclei in nonhuman primates. Journal of Neurosurgery 134, 1064–1071. 10.3171/2019.12.JNS192946

Phillips, K.A., Bales, K.L., Capitanio, J.P., Conley, A., Czoty, P.W., ‘t Hart, B.A., Hopkins, W.D., Hu, S.-L., Miller, L.A., Nader, M.A., Nathanielsz, P.W., Rogers, J., Shively, C.A., Voytko, M.L., 2014. Why primate models matter. American Journal of Primatology 76, 801–827. 10.1002/ajp.22281

Russell, W.M.S., Burch, R.L., 1959. The principles of humane experimental technique. The principles of humane experimental technique.

Sajewski, A., Santini, T., Tiago, M., Berardinelli, J., Ibrahim, T.S., 2023. Comparison of the Optimization of a 60-channel Transmit Coil in pTx and sTx mode at 7T. Presented at the ISMRM.

Santini, T., Wood, S., Krishnamurthy, N., Martins, T., Aizenstein, H.J., Ibrahim, T.S., 2021. Improved 7 Tesla transmit field homogeneity with reduced electromagnetic power deposition using coupled Tic Tac Toe antennas. Sci Rep 11, 3370. 10.1038/s41598-020-79807-9

Santini, T., Zhao, Y., Wood, S., Krishnamurthy, N., Kim, J., Farhat, N., Alkhateeb, S., Martins, T., Koo, M., Zhao, T., Aizenstein, H.J., Ibrahim, T.S., 2018. In-vivo and numerical analysis of the eigenmodes produced by a multi-level Tic-Tac-Toe head transmit array for 7 Tesla MRI. PLOS ONE 13, e0206127. 10.1371/journal.pone.0206127

Saunders, R.C., Aigner, T.G., Frank, J.A., 1990. Magnetic resonance imaging of the rhesus monkey brain: use for stereotactic neurosurgery. Exp Brain Res 81, 443–446. 10.1007/BF00228139

Schaeffer, D.J., Klassen, L.M., Hori, Y., Tian, X., Szczupak, D., Yen, C.C.-C., Cléry, J.C., Gilbert, K.M., Gati, J.S., Menon, R.S., Liu, C., Everling, S., Silva, A.C., 2022. An open access resource for functional brain connectivity from fully awake marmosets. NeuroImage 252, 119030. 10.1016/j.neuroimage.2022.119030

Schneider, C.A., Rasband, W.S., Eliceiri, K.W., 2012. NIH Image to ImageJ: 25 years of image analysis. Nat Methods 9, 671–675. 10.1038/nmeth.2089

Subramanian, T., Deogaonkar, M., Brummer, M., Bakay, R., 2005. MRI guidance improves accuracy of stereotaxic targeting for cell transplantation in parkinsonian monkeys. Experimental Neurology, Including abstracts of the Twelfth Annual Conference of the American Society for Neural Transplantation and Repair 193, 172–180. 10.1016/j.expneurol.2004.11.032

Walbridge, S., Murad, G.J.A., Heiss, J.D., Oldfield, E.H., Lonser, R.R., 2006. Technique for enhanced accuracy and reliability in non-human primate stereotaxy. Journal of Neuroscience Methods 156, 310–313. 10.1016/j.jneumeth.2006.01.025

Xu, F., Jin, H., Yang, X., Sun, X., Wang, Y., Xu, M., Tao, Y., 2018. Improved accuracy using a modified registration method of ROSA in deep brain stimulation surgery. Neurosurgical Focus 45, E18. 10.3171/2018.4.FOCUS1815

Yazdan-Shahmorad, A., Tian, N., Kharazia, V., Samaranch, L., Kells, A., Bringas, J., He, J., Bankiewicz, K., Sabes, P.N., 2018. Widespread optogenetic expression in macaque cortex obtained with MR-guided, convection enhanced delivery (CED) of AAV vector to the thalamus. Journal of Neuroscience Methods 293, 347–358. 10.1016/j.jneumeth.2017.10.009

Zhu, G.-Y., Chen, Y.-C., Du, T.-T., Liu, D.-F., Zhang, X., Liu, Y.-Y., Yuan, T.-S., Shi, L., Zhang, J.-G., 2019. The Accuracy and Feasibility of Robotic Assisted Lead Implantation in Nonhuman Primates. Neuromodulation: Technology at the Neural Interface 22, 441–450. 10.1111/ner.12951

